# Super-resolution imaging reveals dynamic reticular cytoophidia

**DOI:** 10.1101/2022.08.29.505636

**Authors:** Yifan Fang, Yi-Lan Li, Xiao-Ming Li, Ji-Long Liu

**Affiliations:** School of Life Science and Technology, ShanghaiTech University, Shanghai, China; Department of Physiology, Anatomy and Genetics, University of Oxford, Oxford, OX1 3PT, United Kingdom

**Keywords:** CTP synthase, cytoophidium, fluorescence recovery after photobleaching (FRAP), Stimulated emission depletion (STED)

## Abstract

CTP synthase (CTPS) can form filamentous structures termed cytoophidia in cells from all three domains of life. In order to study the mesoscale structure of cytoophidia, we perform fluorescence recovery after photobleaching (FRAP) and stimulated emission depletion (STED) microscopy in human cells. By using EGFP dimeric tag, as a tool to explore the physical properties of cytoophidia, we find that cytoophidia are dynamic and reticular. The reticular structure of CTPS cytoophidia may provide space for other components such as IMPDH. In addition, we observe CTPS granules with tentacles.

## 1. Introduction

In addition to organelles with membrane, proteins with important functions in the cell can also be compartmented into membraneless organelles. Protein compartmentation, like droplets, can be assembled driven by physical forces in cells (Brangwynne et al., 2009). CTP synthase (CTPS), a metabolic enzyme for *de novo* synthesis of CTP, was found to form filament-like compartments in cells called cytoophidia (Ingerson-Mahar et al., 2010; Liu, 2010; Noree et al., 2010). For describing the shape vividly, cytoophidia mean cellular snakes in Greek. Cytoophidia are conserved in evolution (Liu, 2016).

A glutamine analog, 6-diazo-5-oxo-L-norleucine (DON), promotes cytoophidia formation in *Drosophila* and human cells (Chen et al., 2011). DON binds CTPS with covalent bonds (Zhou et al., 2021). Glutamine deprivation promotes cytoophidium formation in mammalian cells (Calise et al., 2014). IMPDH can form cytoophidia (Ji et al., 2006) and both CTPS and IMPDH are related to glutamine and NH3 metabolism. It was reported that there is an interaction between CTPS and IMPDH (Chang et al., 2018). The function of cytoophidia may be closely related to glutamine and NH3 metabolism.

In metabolic regulation, the activity of CTPS is inhibited via filament formation (Barry et al., 2014; Lynch et al., 2017). The half-life of CTPS is prolonged when forming cytoophidia (Sun and Liu, 2019a). With high-level metabolism in cancer cells, cytoophidium formation is highly related to oncogenes. Myc is required for cytoophidia assembly and cytoophidia formation is regulated by Myc expression level (Aughey et al., 2016). Ack kinase regulates cytoophidium morphology and CTPS activity (Strochlic et al., 2014). Cytoophidium assembly was regulated by mTOR-S6K1 pathway (Sun and Liu, 2019b). Cytoophidia were found in human hepatocellular carcinoma (Chang et al., 2017).

CTPS can be assemble into thin filaments in vitro, and the structures of CTPS filaments at near-atomic resolution have been solved by cryo-EM (Lynch et al., 2017; Zhou et al., 2019). However, nanometer scale CTPS filaments are different from micron scale cytoophidia observed under confocal microscopy. How CTPS filaments assembles into big micron scale cytoophidia is still unclear. The physical properties of cytoophidia at the mesoscale remain to be explored.

To study cytoophidium properties in human cell lines, we performed fluorescence recovery after photobleaching (FRAP) microscopy to study the dynamic characteristics and stimulated emission depletion (STED) microscopy to study the super-resolution structure. By measuring the intensity and recovery speed of bleaching ROI, we can quantify the relative dynamic characteristics of cytoophidia with different treatments. STED allows fluorescence imaging to achieve a resolution of 50 to 70 nm (Jans et al., 2013; Sezgin et al., 2017).

## 2. Results

### 2.1. Assembly of CTPS filaments into cytoophidia

The cytoophidium is a compartment of metabolic enzymes, such as filamentous CTPS, which can be observed under confocal microscopy (Ingerson-Mahar et al., 2010; Liu, 2010; Noree et al., 2010). In vitro experiments showed that CTPS can also form filaments building from tetramer units (Lynch et al., 2017; Zhou et al., 2019). In human cells, CTPS can be assembled into cytoophidia under DON treatment or glutamine deprivation. In this study, we did not distinguish hCTPS1 and hCTPS2. We found that after DON treatment, hCTPS1 can form granules in 293T cells with hCTPS1 over-expression (Figure 1A and B). When we observed living cells, CTPS granules existed in a small population of cells. CTPS granules can exist in the same cell with cytoophidia (Figure 1A-C).

**Figure 1.**
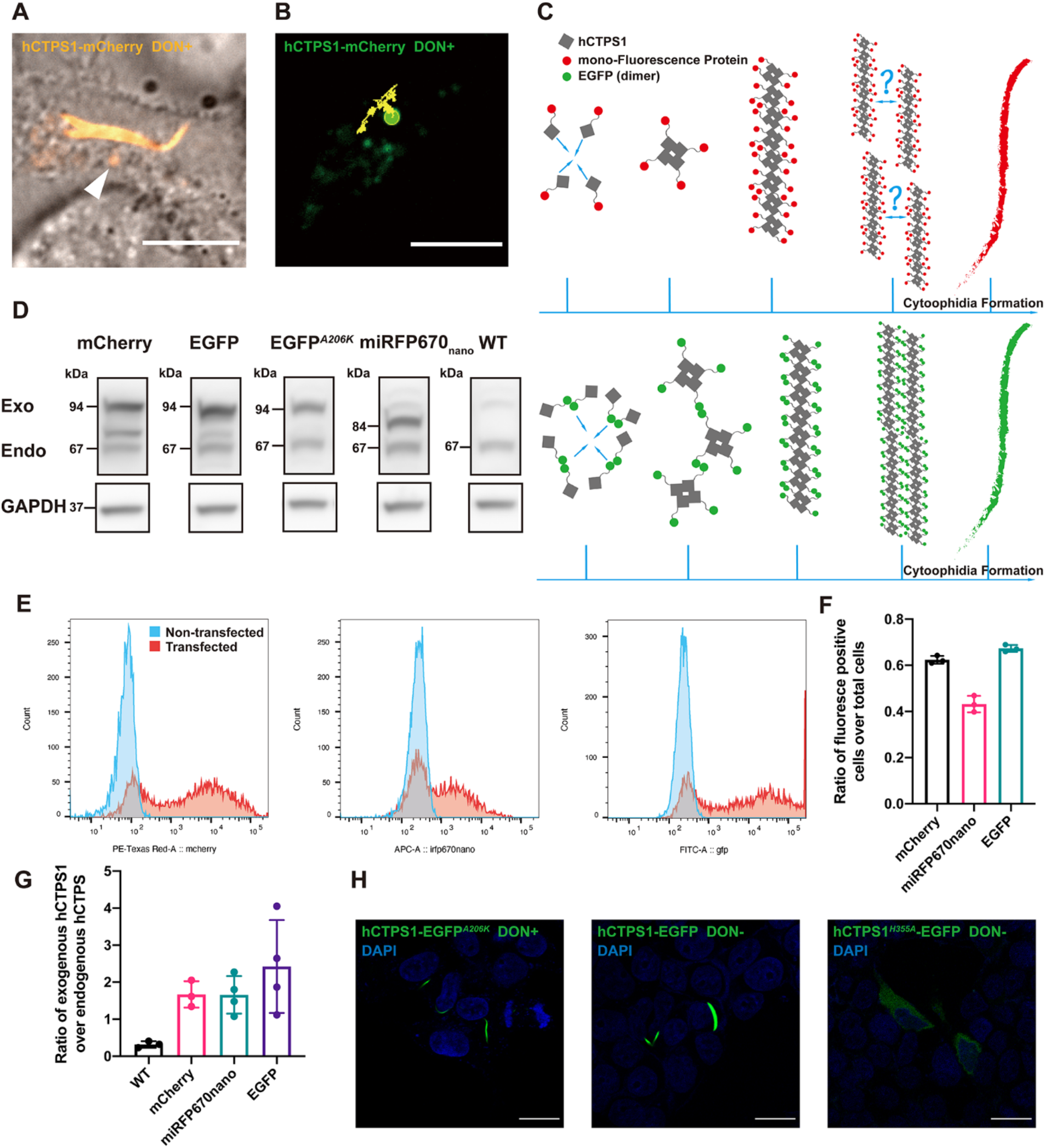
Assembly of CTPS filaments into cytoophidia. (**A**) hCTPS1 forms granules in the same cell that coexist with hCTPS1 cytoophidia. The arrow head points out the granule (**B**) The trajectory of hCTPS1 granules is a random walk. For DON treatment, 20 µg/mL DON in PBS solution was added into fresh DMEM media 8 to 25 hours before living cell imaging. (**C**) hCTPS1-EGFP forms cytoophidium-like condensates by simple force between EGFP. The arrangement from hCTPS1 filament to hCTPS cytoophidia is the problem that need to be solved. (**D**) The quantities of transfected over-expression hCTPS in 293T cells were measured respectively. (**E** and **F**) The transfection efficiencies were quantified respectively. (**G**) Estimated ratio of exogenous hCTPS1 to endogenous hCTPS. (**H**) hCTPS-EGFPA206K cytoophidia can be induced by DON treatment. hCTPS1-EGFP can form cytoophidium-like condensates, which is wider and larger than hCTPS cytoophidium. hCTPS1H355A-EGFP cannot form cytoophidium-like condensates. For DON treatment, 20 µg/mL DON (PBS solution) was added into fresh DMEM media 8 hours before fixation. Scale bars, 10 µm (**A, B**) and 20 µm (**H**).

How CTPS filaments are arranged in cytoophidia remains unclear. To conceive the arrangement model from CTPS filaments to cytoophidia, we constructed hCTPS1 overexpression vectors with different fluorescence proteins and different mutations (Supplemental Table S1). In order to show and evaluate the effect of exogenous overexpression protein on the experiment, we compared the protein levels of overexpressed hCTPS1 and endogenous hCTPS1/2 (Figure 1D), and tested the transfection efficiencies (Figure 1E and F). In transfection positive cells, exogenous hCTPS1 was approximately twice as much as endogenous hCTPS1/2 (Figure 1G).

EGFP is a weak dimer (Shaner et al., 2005). Overexpression of hCTPS1-EGFP forms cytoophidium-like condensates in 293T cells (Figure 1H). The force of forming dimer between EGFP pulls hCTPS1 together, and hCTPS is assembled into filaments. The hCTPS filaments may be compressed together by a simple force of EGFP dimerization(Figure 1C). The hCTPS1-EGFP group was a control for cytoophidium induction. CTPS1 with H355A mutation disassembles cytoophidia. We found that overexpressed hCTPS1^*H355A*^-EGFP could not form cytoophidia in 293T cells (Figure 1H).

### 2.2. Dynamic equilibrium of cytoophidia

To test the dynamic characteristics of cytoophidia, we performed FRAP on four groups of hCTPS1 cytoophidia (Figure 2A and B). The intensity of FRAP ROI was normalized as 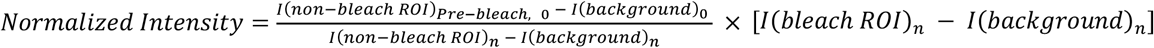 (Warrington et al., 2022). The bleached ROI on cytoophidia induced by 20 µg/mL for 8 hours (low concentration and short time) before imaging recovered very quickly (Figure 2C). However, ROI in cells with hCTPS1 overexpression treated with DON in 100 µg/mL (higher concentration) recovered fluorescence slowly and ended at a lower intensity (Figure 2D; Supplemental Figure S1A).

**Figure 2.**
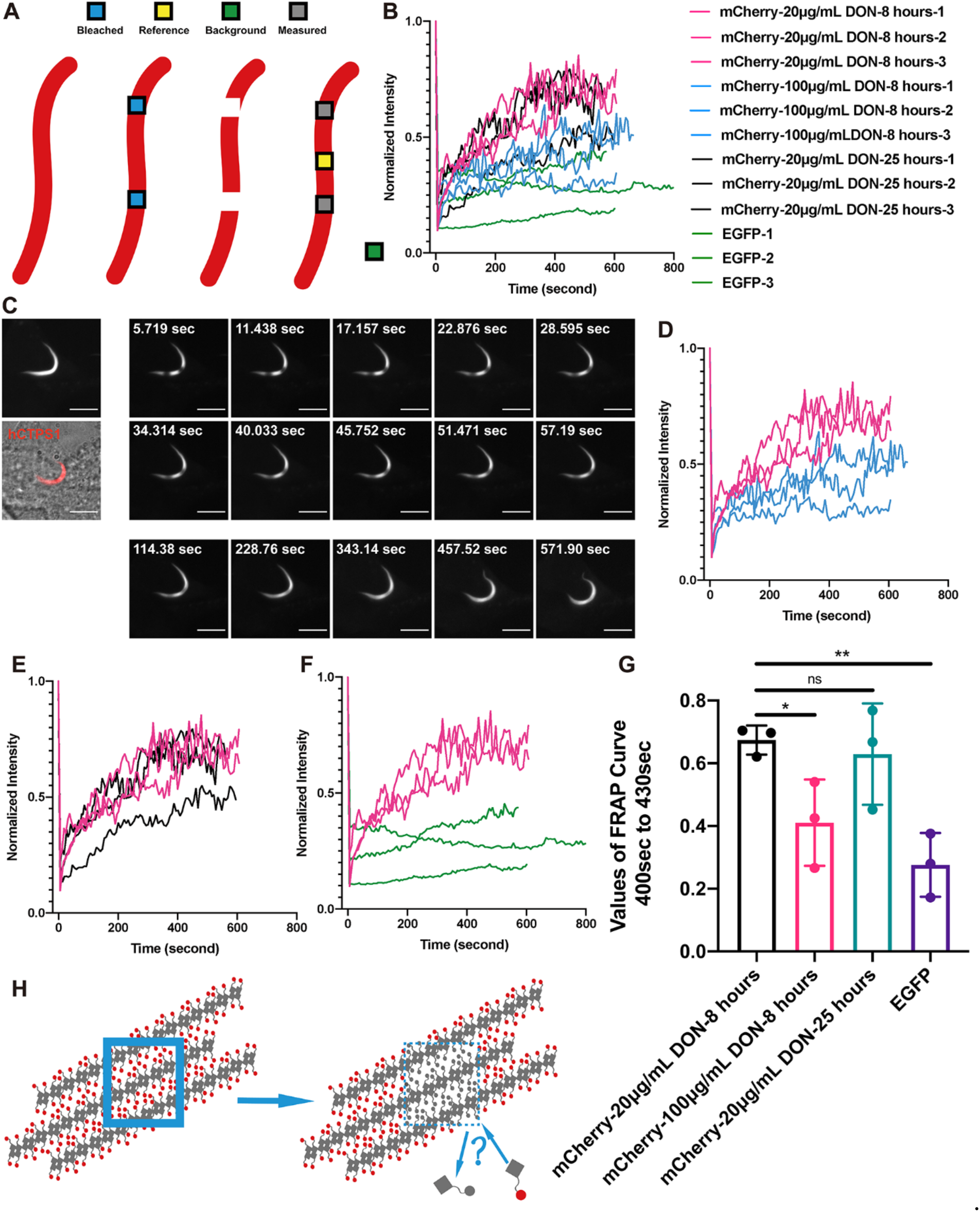
Dynamic equilibrium of cytoophidia. (**A**) ROIs were used for bleaching, measurement and standardization data. (**B**) Normalized intensity curves of FRAP results in different groups were merged. (**C**) Living cell images of FRAP on hCTPS1-mCherry cytoophidia induced by 20 µg/mL DON for 8 hours. (**D**) Comparison of FRAP curve on hCTPS1-mCherry between 20 µg/mL DON for 8 hours and 100 µg/mL DON for 8 hours. (**E**) Comparison of FRAP curve on hCTPS1-mCherry between 20 µg/mL DON for 8 hours (pink curve) and 20 µg/mL DON for 25 hours (black curve). (**F**) Comparison of FRAP curve between hCTPS1-mCherry cytoophidia (pink curve) induced by DON and hCTPS1-EGFP cytoophidium-like condensates (green curve). (**G**) Analysis of the level and speed of fluorescence recovery *, p-value<0.05; **, p-value<0.01; ns, no significant difference. (**H**) It does not fit the results of FRAP images if hCTPS cytoophidia are compact or condensed. A new model is needed to illustrate the recovery of bleached fluorescence of cytoophidia. The intensity of FRAP ROI was normalized as 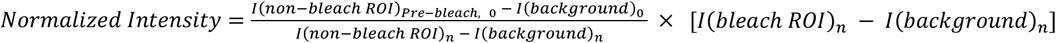. Scale bars, 10 µm (C).

By extending the DON treatment time to 25 hours, ROI can recover as soon as possible in a relatively short time (Figure 2E; Supplemental Figure S1B). The fluorescence intensity restored by bleaching ROI of cytoophidium-like condensates of hCTPS-EGFP cells is very small (Figure 2F; Supplemental Figure S1C)., There was a significant difference between hCTPS1-EGFP cytoophidium-like condensates and DON-induced hCTPS cytoophidia at low concentration and in a short time (Figure 2G).

This shows that DON-induced cytoophidia have very different dynamic characteristics from hCTPS-EGFP cytoophidium-like condensates. DON-induced cytoophidia seem not assembled by simple force like hCTPS-EGFP cytoophidium-like condensates.

The bleached ROI gradually recovered throughout the ROI, rather than from any side of the ROI. Neither of the two ROIs moved to either side, nor were they far away or close to each other (Figure 2C). We observed that bleached hCTPS1 molecules in cytoophidia could exchange with free hCTPS1 molecules in the cytosol (Figure 2H).

### 2.3. The reticular structure of hCTPS1 cytoophidium and its localization with hIMPDH2

In order to build a model to fit the dynamic equilibrium characteristics of cytoophidia, we obtained the super-resolution structures of hCTPS cytoophidia using stimulated emission depletion microscopy (STED). The images under conventional confocal microscopy could not show the structure inside cytoophidia, while the STED images revealed the super-resolution structure with a resolution of 50 to 70 nm (Figure 3A; Figure S3A, B), which implied a possible mechanism of highly dynamic cytoophidia under FRAP. We estimated the resolution by measuring the distance between two distinguishable nearby particles and it was 50 to 70 nm (Figure S3A, B).

**Figure 3.**
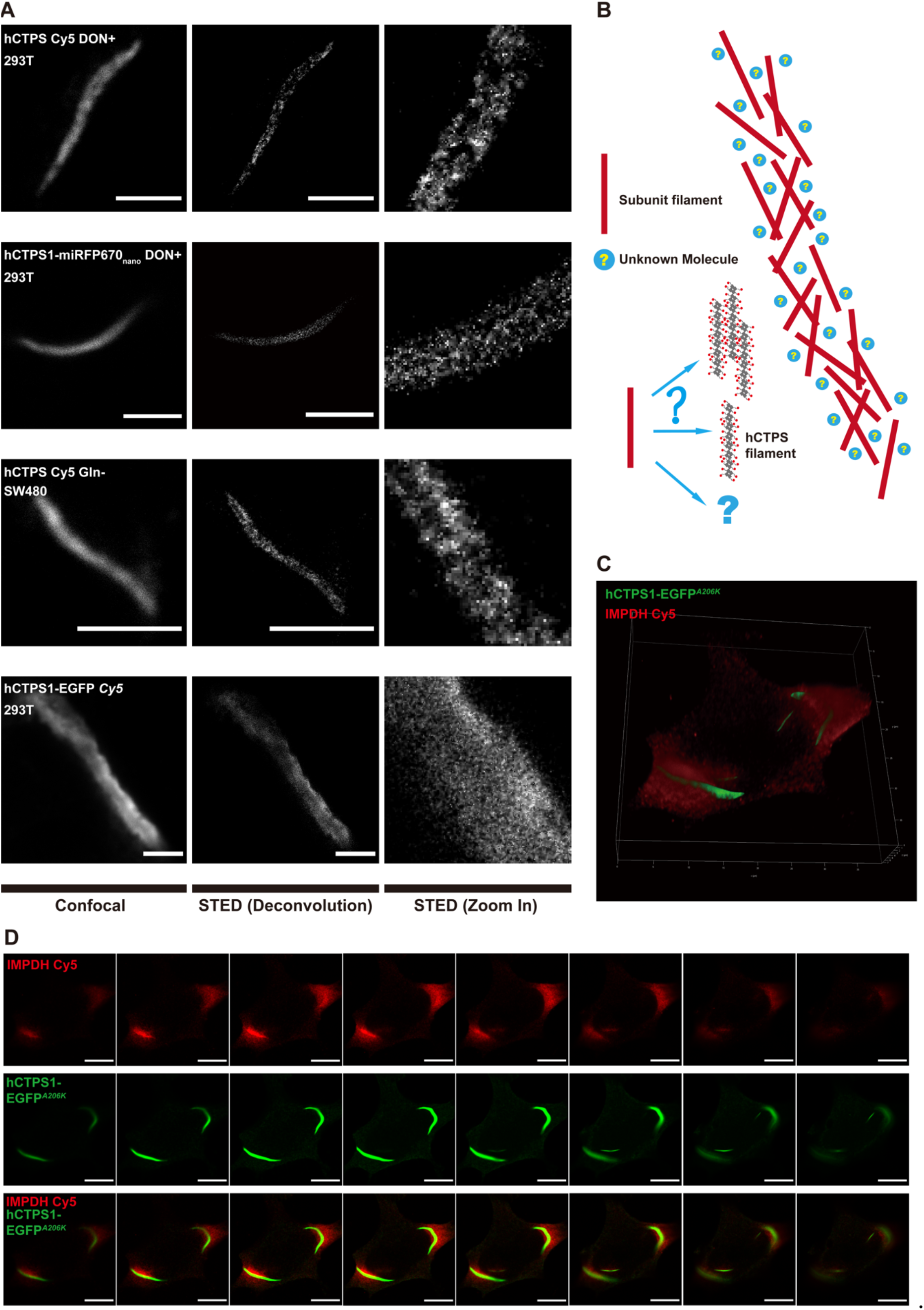
The reticular structure of hCTPS1 cytoophidium and its localization with hIMPDH2. (**A**) Confocal, STED with deconvolution and zoom-in images of cytoophidia from the following groups: 1) hCTPS1/2 cytoophidia (Cy5 antibody stained) induced by DON in fixed 293T cells, 2) hCTPS1-miRFP670nano cytoophidia induced by DON in live 293T cells, 3) hCTPS1/2 (Cy5 stained) cytoophidia induced by glutamine deprivation in fixed SW480 cell and 4) hCTPS1/2 (Cy5 stained)-cytoophidium-like condensates in live 293T (hCTPS1-EGFP overexpression) cell. For SW480 culture, DMEM without glutamine replaced DMEM 8 hours before fixed. For DON treatment, DON (PBS solution) was added into fresh DMEM media 8 to 25 hours before living cell imaging or 8 hours before being fixed. Scale bars, 3 µm. (**B**) The arrangement model of hCTPS filaments into cytoophidia. (**C**) In 293T cells, hCTPS1-EGFPA206K was colocalized with DON induced IMPDH2(Cy5 stained). (**D**) In each slice along Z stacks, hCTPS1-EGFPA206K and IMPDH2 (Cy5 stained) are localized to each other. Scale bars, 10 µm.

STED revealed a heterogeneous structure of hCTPS cytoophidia. Some parts of it were more condensed, while other parts were looser. More importantly, it seemed that there were many tiny filaments inside, which in different orientations constructed into a reticular structure (Figure 3A; Supplemental Figure S2A).

However, the super-resolution results were homogeneous inside hCTPS1-EGFP cytoophidium-like condensates (Figure 3A). It was totally different between the structure of hCTPS cytoophidia and that of hCTPS1-EGFP. No condensed and loose difference and reticular structure knitted with tiny filaments can be observed in cytoophidium-like condensates.

When doing super-resolution imaging, there might be some influencing factors, such as the efficiency of antibodies, optical properties of fluorescent labeling, the steric hindrance of fluorescence proteins, and the influence of sample preparation. To eliminate the effects of antibodies and Cy5, we used 293T cells with hCTPS1-miRFP670nano for live cell imaging. We performed immunofluorescence staining on DON-treated 293T cells to eliminate the effects of steric hindrance and overexpression. Both results showed reticular structures (Figure 3A). To avoid the difference of optical properties between Cy5 and EGFP, we also performed immunofluorescence staining with Cy5 on hCTPS1-EGFP over-expressed 293T cells and signal obtained was from Cy5 (Figure 3A).

In addition, we performed immunofluorescence staining on SW480 cells cultured in glutamine free medium, which showed a reticular structure (Figure 3A). Glutamine is the NH3 donor in metabolic reaction. This means that the reticular structure of hCTPS is not only a phenomenon induced by DON, but also a common structure of metabolic enzymes when cells are under metabolic stress. Without changing the super-resolution structure results, deconvolution of the STED images could improve their resolution and signal-to-noise ratio (Figure 3A; Supplemental Figure S2A).

These tiny filaments appear as subunits of hCTPS cytoophidia (Figure 3A and B). In vitro experiments show that CTPS can be assembled into filaments (Lynch et al., 2017; Zhou et al., 2019). Based on the in vitro and in vivo results, we envisioned a model to illustrate the reticular structure of hCTPS cytoophidia (Figure 3B). Inside cytoophidia, subunit filaments weaved into a reticulation. The model can make FRAP results clearer. The dynamic equilibrium of assembly and disassembly occurs in the tiny filaments of hCTPS, rather than the assembly and disassembly of the whole cytoophidia. FRAP on untreated and dispersive hCTPS1 signal resulted in fast recovery, which made us unable to capture the image after bleaching, which was similar to the image before bleaching (Supplemental Figure S2B). Because of the limitation of STED resolution, it is unclear that the subunit filament is one CTPS filament, a bundle of CTPS filaments or some other form of CTPS.

After obtaining the super-resolution reticular structures, we wanted to know the reason and function of this reticular structure. There might be some unknown molecules in the space between hCTPS filaments (Figure 3B). It was reported that IMPDH2 interacted with CTPS1 cytoophidia under DON treatment (Chang et al., 2018). Cytoophidia of hCTPS1 and hCTPS2 located together in 293T cells (Supplemental Figure S2C). IMPDH and CTPS are both part of the glutamine and NH3 metabolic pathways (Supplemental Figure S2D). We overexpressed hCTPS1-EGFP^*A206K*^, a monomer version of EGFP, and labeled IMPDH2 with Cy5 by immunofluorescence staining. Under DON treatment, IMPDH2 and hCTPS1 were colocalized spatially (Figure 3C). IMPDH2 and hCTPS1 were not exclusive, but were positioned mutually in each Z stack (Figure 3D). Therefore, IMPDH2 could be one of the molecules located between hCTPS1 filaments.

### 2.4. CTPS granules with tentacles

We found that hCTPS1 could not only form cytoophidia, but also formed DON induced granules (Figure 1A). The movement of most granules was random walk like that of ordinary granules in cells (Figure 1B). Fortunately, when we observed live cells treated with DON, we found a filiform structure connected between hCTPS1 granules (Figure 4A). We name these filiform structures connecting granules “tentacles”, and the main part of the granules is called the “granule body”. The tentacle slowly extended out of the granules body and retracted as quickly as a rubber band after reaching its longest length (Figure 4B).

**Figure 4.**
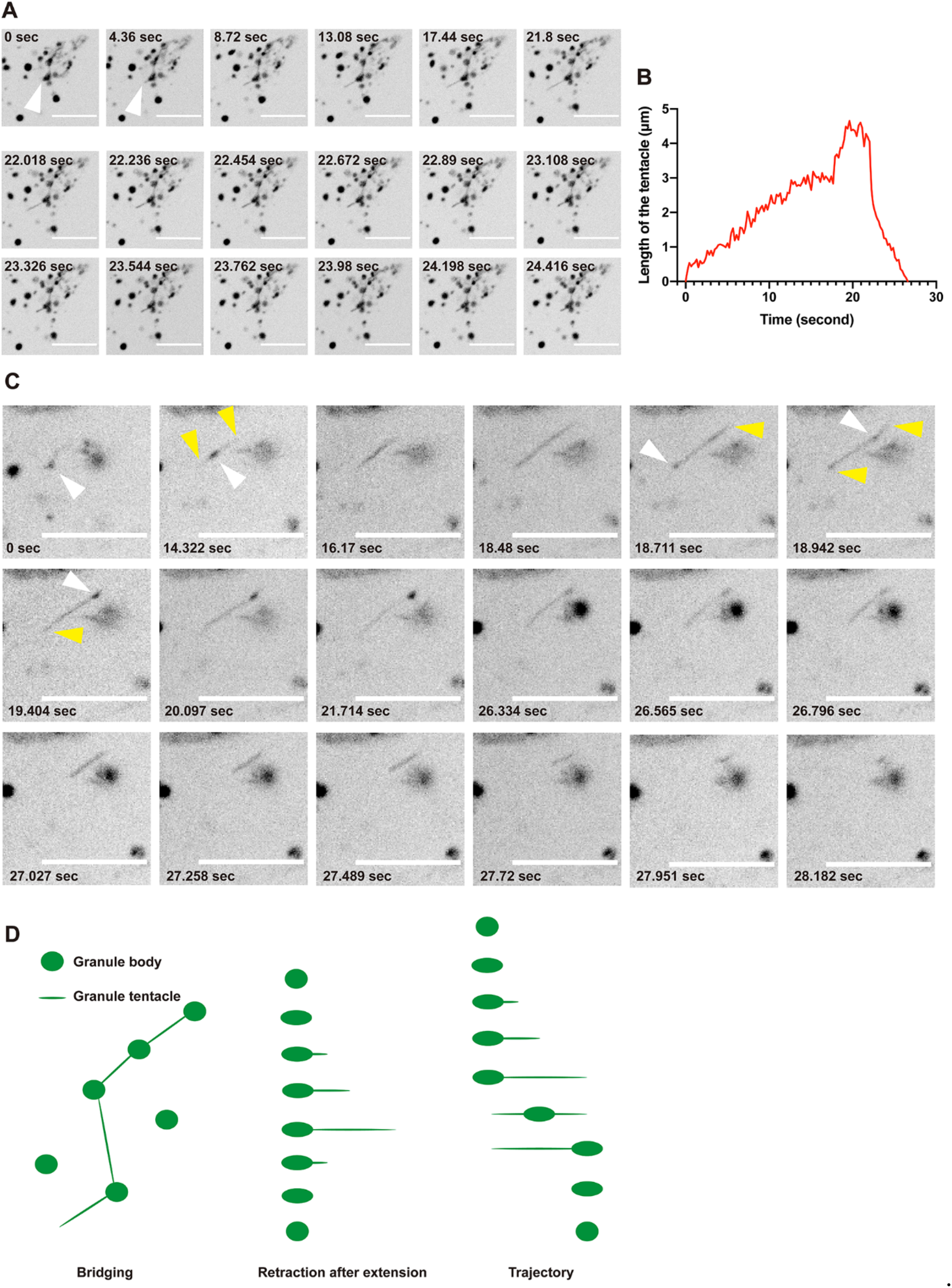
CTPS granules with tentacles. (**A**) Tentacles connect hCTPS1 granules. Tentacles extend and retract. Arrow head indicates the tentacle. (**B**) Tentacles extend slowly and retract rapidly after reaching the maximum length. (**C**) hCTPS1 granules move from one side to the other along the tentacles. Yellow arrow heads indicate the tentacles. White arrow heads indicate the granules. (**D**) hCTPS1 granule tentacles have three different behaviors and characteristics, bridging, retraction after extension, and the trajectory of hCTPS1 granule movement. For DON treatment, DON (PBS solution) was added to fresh DMEM media 8 to 25 hours before living cell imaging. Scale bars, 10 µm (**A, C**).

We wanted to know the function of the granule tentacles. We found that tentacles were different from small granule bodies. When the tentacle stretched out, the granule body moved from one side of tentacle to the other side along the tentacle, and then tentacle retracted the granule body in the new location (Figure 4C). Granules with tentacle move with clear direction along the tentacles rather than random walk. The movement of tentacle granules was different from that of non-tentacle granules. Granular tentacles are tiny structures that bridge granules, move granules and retract after extension (Figure 4D).

## 3. Discussion

Taking advantage of multiple fluorescence tags, we study the physical characteristics of hCTPS1-containing compartments under light microscopy. We perform FRAP and STED to reveal the dynamic and reticular structure of cytoophidia. In addition, we observe that hCTPS1 forms granules with tentacles.

### 3.1. Cytoophidia are not condensates

Since the discovery of CTPS forming cytoophidia, the exact phase of cytoophidia and the arrangement of CTPS in cytoophidia are still unclear. Cytoophidia are presumed to be static bundles of filaments (Liu, 2011) or liquid phase, just like LLPS. However, when we performed live cell imaging on CTPS cytoophidia, we found that CTPS can not only form long filamentous structures, that is, cytoophidia, but also form granules in the same cell (Figure 1A). This means that as a compartment of CTPS, cytoophidia may not be a static, concentrated and rigid structure. In another hypothesis, the puzzling question is why this compartment is not a spherical droplet if it is in the liquid phase, such as LLPS.

According to previous studies, the residue 355H of CTPS (CTPS-355H) is the key site to form this filament structure. CTPS-355H lying at the tetramer-tetramer interface plays a critical role in CTPS polymerization. In vitro experiments, the CTPS tetramer assembly mechanism of cytoophidia is more like actin filament than droplets in cells assembled by physical force (Banani et al., 2017; Zhang, 2020).

To study the role of CTPS-355H, we used both dimeric EGFP and monomeric EGFP^*A206K*^ tags (Shaner et al., 2005). We generated hCTPS1^*H355A*^ mutations. hCTPS1-EGFP^*A206K*^ can form cytoophidia with DON treatment. Without DON treatment, hCTPS1-EGFP^*A206K*^ cannot form cytoophidia, suggesting EGFP^*A206K*^ does not promote CTPS assembling. mCherry and miRFP670nano are also monomeric tags just like EGFP^*A206K*^. However, hCTPS1-EGFP can form filament-shaped condensates without DON treatment (Figure S2E). Because EGFP has a force to form dimer like ‘sticky’ feature, and hCTPS1-EGFP stick to each other into filament-shaped condensates, we refer these filament-shaped hCTPS1-EGFP structures as ‘cytoophidium-like condensates’ (to be distinguished from the term ‘cytoophidia’) (Figure 1C).

Are cytoophidia just condensates? If this is true, the key cytoophidium-forming site CTPS-355H may provide a directional force, which is important for condensate formation. For hCTPS1^*H355A*^-EGFP, the directional force is provided by dimeric EGFP, since CTPS-355H has been mutated to CTPS^*H355A*^. If either CTPS-355H or EGFP can provide directional force for condensate formation, we would expect that both hCTPS1^*H355A*^-EGFP and hCTPS1-EGFP^*A206K*^ can form condensates. Our results show that hCTPS1^*H355A*^-EGFP cannot form cytoophidium-like condensates, suggesting that CTPS-H355 is an essential site of connection rather than just providing a directional force (Figure 1H).

Therefore, our data argue against the idea that cytoophidia are condensates. Two factors appear to be required for assembling CTPS into cytoophidia. The first, CTPS molecules are bring together by some forces of assembling. The second, CTPS tetramers need to be connected via CTPS-355H.

### 3.2. Cytoophidia are dynamic

In order to solve the problem of the physical phase of cytoophidia, we carried out FRAP to measure the dynamic features. We used hCTPS1^*H355A*^-mCherry as the control of complete diffusion, which recovered its intensity quickly after bleaching, so that we could not capture the difference before and after bleaching or measure its dynamic value. We used hCTPS1-EGFP cytoophidium-like condensates as a static control. The intensity recovered from cytoophidia was significantly faster than that of hCTPS1-EGFP cytoophidium-like condensates (Figure 2F, G). This means that the cytoophidium is not a condensate, but a highly dynamic structure.

Low concentration of DON can induce cytoophidia, mimicking the stress of severe glutamine deprivation. We also measured the effects of time and concentration of DON treatment to determine whether the dynamic results were valid only in particular circumstances. Compared with these curves, the treatment time of DON had little effect on the kinetics (Figure 2E, G), while the concentration of DON had a significant effect on the kenetics of cytoophidia (Figure 2D, G). Due to the covalent bond between DON and CTPS, excessive DON may destroy the conformation of CTPS, thus damaging cells.

### 3.3. Cytoophidia are reticular

The cytoophidium is micrometer scale. A previous assumption was that the cytoophidium might be a bundle of CTPS filaments, similar to actin filaments. When the bleached ROI gradually recovered its intensity, no treadmill phenomenon was observed, which means that the assembly mechanism of cytoophidia is different from that of actin filaments. Moreover, recovery does not come from either side of the bleached ROI. The intensity of the entire ROI recovers steadily at the same speed.

If the cytoophidium is a bundle of CTPS filaments, how can the bleached ROI recover its intensity without treadmill phenomenon (Figure 2H)? To solve this problem, we need to solve the fine structure of cytoophidia. We performed super-resolution STED imaging which revealed the reticular structure of cytoophidia. In order to minimize the artificial influence on the imaging results, we used miRFP670nano as a fluorescence tag to conduct STED imaging directly on live cell samples, which gives a clear reticular structure (Figure 3A). We call it reticular structure because small subunit filaments are interconnected, crossed and woven into reticulation (Figure 3B).

At present, it is unclear whether the subunit filament is a CTPS filament, a bundle of CTPS filaments or other forms of CTPS. To maximize the resolution of STED imaging (Fig. 3B), all fluorescent tags are infrared emitting. miRFP670nano is much smaller than other infrared fluorescent proteins, and it can minimize the impact of tag space obstruction. We performed immunofluorescence staining to confirm the results and avoid the effects of spatial obstruction and overexpression. Cytoophidia with immunofluorescence dye Cy5 on hCTPS also shows the reticular structure.

We also performed immunofluorescence staining on SW480 cells. Cytoophidia in SW480 cells can be induced by glutamine deprivation and present reticular structures, which means reticular cytoophidia are common in different cell types and under glutamine metabolic stress condition, but cannot be treated artificially. Even DON is used on cells to mimic the stress of glutamine metabolic stress.

We used hCTPS1-EGFP as the structurally static control to represent the condensate assembled by physical forces. hCTPS1-EGFP cytoophidium-like condensates do not show the reticular characteristics. The cytoophidium-like condensates appear homogeneous, and their internal parts look the same. There is neither subunit filaments nor space can be observed inside cytoophidium-like condensates. In summary, the structure of cytoophidium is reticular, and the arrangement of CTPS assembled on cytoophidia is different from that of condensates or actin filaments.

The cytoophidium reticulation provides a structural basis for the localization of other enzymes such as IMPDH (Figure 3C, D). Both CTPS and IMPDH are associated with glutamine and NH3 metabolism. In reticular cytoophidia, there may be dynamic interaction between CTPS, IMPDH and their substrates and the microenvironment. The reticular structure can also provide elasticity for the bending or twisting of cytoophidia (Figure S1B). It was reported that cytoophidia are related to the regulation of IMPDH activity (Chang et al., 2015).

The reticular structure provides a structural basis for FRAP results. No treadmill phenomenon has been observed. How can CTPS molecules in free state replace those composed the bleached ROIs? Based on the assumption of the reticular structure, the treadmill may occur in the subunit filaments rather than the whole cytoophidia. The assembly and disassembly of CTPS on subunit filaments may have a dynamic equilibrium, thus changing the CTPS molecules after bleaching. This may be a potential explanation for the dynamic equilibrium of large scale FRAP.

Due to the limitations of STED or confocal devices, we cannot achieve super-resolution, high-speed live cell imaging and low phototoxicity to cells. In order to verify this speculation, more advanced microscope technology is required. Dynamic equilibrium of assembly and disassembly in subunit filaments may contribute to metabolic regulation of reactions in the microenvironment.

In *Cautobacter crescentus*, a small amount of CTPS forms a bundle, while a large amount of CTPS forms a splayed structure (Ingerson-Mahar et al., 2010). A large number of hCTPS transforms the morphology of CTPS bundles into complex structures. Cytoophidia in *Drosophila* female germ cells also exhibit reticular characteristics (Liu, 2010). Cytoophidia are regulated by the level of molecular crowding in the cell (Chang et al., 2022). The formation and maintenance of the reticular cytoophidia may be related to molecular crowding.

### 3.4. CTPS can form granules with tentacles

While using live cell imaging to capture CTPS-containing structures, we found interesting CTPS granules with tentacles. CTPS granules move in a random walking mode, but the granules with tentacles move in a clear direction. We assume that these two are different forms of compartments with similar shapes. We call it the tentacle because it slowly stretches out of the granules and quickly retracts, just like the tentacle of an octopus or snail (Figure 4B). The tentacle may be extended to find something that can be connected. The tentacles, like bridges, connect different granules (Figure 4A).

If the granules are very small, the tentacles play a role in directional movement (Figure 4C). The granules extend the tentacles to the maximal length, and the granules move quickly from one side to the other along the tentacles, just like a slingshot. The only function of tentacles we know is related to the directional movement of granules. We still do not know the function of tentacles as bridges. Moreover, it is unclear whether the granules with tentacles have membrane, because this movement is similar to the movement of mitochondria or the vesicles from Golgi.

Interestingly, CTPS granules, tentacled granules and cytoophidia appeared under the same conditions, i.e. treated with DON. They can even exist in the same cell, with potential interaction and transformation. The tentacles also have directions, just like cytoophidia. However, due to the tiny size, the tentacle may not share the same reticular structure as the cytoophidium.

The structure of the tentacle may be similar to the subunit CTPS filament to obtain directional characteristics, or it may be in the liquid state in the membrane vesicle. The intensity of granules with tentacles is far lower than that of cytoophidia, and the tentacles exist in fewer cells than cytoophidia do. Because of its low intensity and small volume, it is difficult to analyze the properties and fine structures of the tentacle. Compared with cytoophidia reacting to metabolism, it is not clear whether granules with tentacles are related to glutamine metabolism or the role of DON.

### 3.5. Conclusion

To sum up, the main purpose of this study is to understand the structure and arrangement of CTPS between CTPS filament with near-atomic resolution structure and micron scale cytoophidia observed under confocal microscopy. We use dimeric EGFP tag as a control to provide aggregation viscosity, and identify the connecting role of the CTPS-355H site. FRAP analysis shows that the cytoophidium is highly dynamic, while STED analysis reveals the reticular structure of cytoophidia.

According to the comparison with CTPS-EGFP cytoophidium-like condensates, the dynamic and reticular characteristics of cytoophidia are different from those of condensates (Figure 3B). Moreover, we find that the compartment of CTPS not only exists in the snake-shaped cytoophidia, but also exists in the granules. CTPS granules move in different ways depending on whether they have tentacles. Functions of CTPS granules with tentacles require further studies.

## 4. Materials and Methods

### 4.1. Cell culture

293T and SW480 were cultured in DMEM (SH30022.01, Hyclone) supplemented with 10% FBS (04–001; Biological Industries) in a humidified atmosphere containing 5% CO2 at 37°C. All the commercial cell lines used in this article were purchased from Shanghai Institutes for Biological Sciences, Chinese Academy of Sciences (Shanghai, China). They were originally purchased from ATCC. DON was dissolved in PBS. DON were added to cultured medium as described in individual experiments. DMEM without glutamine (C11960500BT, Gibco) replaced DMEM 8 hours before imaging.

### 4.2. Constructs and transfection

The pLV-hCTPS1-EGFP over-expression vector was kindly provided by Dr. Zhe Sun from ShanghaiTech University. mCherry, miRFP670nano replaced EGFP and EGFP^*A206K*^ mutated back into EGFP using PCR and Gibson Assembly System (NEB). Cell transfection was done with PEI reagent (24765-1, Polysciences) according to instructions provided by the manufacturers. The sequences of oligonucleotides used in this study are listed in Supplemental Table S2.

### 4.3. Immunoblotting

Cells were harvested in lysis buffer (Containing 20mM Tris, 150mM NaCl and 1% Triton X-100; P0013J, Beyotime). Undissolved cell fraction was separated by centrifugation at 12000 rpm for 10 min at 4°C and the supernatant were boiled in SDS-PAGE loading buffer for 10 min. Proteins in total cell lysates were separated by SDS-PAGE and transferred to PVDF membranes. Membranes were blocked in 5% nonfat milk and incubated with the appropriate primary antibodies. Protein bands were visualized through horseradish peroxidase (HRP) conjugated secondary antibody by ECL reagent (34577, ThermoFisher Scientific).

### 4.4. Immunofluorescence

Cells were fixed with 4% paraformaldehyde added into media for 25 min. Then the fixed cells were washed in 1xPBS for 3 times. Samples were incubated with appropriate primary antibodies (rabbit anti IMPDH2, Proteintech 12948-1-AP; rabbit anti CTPS, Proteintech 15914-1-AP) overnight at 4 °C, washed in PBS for 3 times. Samples were incubated with Cy5 conjugated secondary antibodies (donkey anti rabbit Cy5 conjugated antibody, Jackson 711-175-152) at room temperature for 1 hour (in dark), and washed with PBS for 3 times after incubation. Mountant for STED imaging (Figure 3A, C, D; Figure S2A) was Prolong™ Diamond Antifade (Invitrogen. P36965). Mountant for confocal imaging (Figure 1H) was HardSet Mounting Medium with DAPI (VECTASHIELD, H-1500)

### 4.5. Microscopy

Images (Figure 1A, B; Figure 2C; Figure 4A, C; Figure S1A, B, C; Figure S2B) were acquired under 100× objectives on a confocal microscope (Nikon CSU-W1 SoRa). Confocal images and super-resolution images (Figure 3A, C, D; Figure S2A) were acquired under 100× objectives on a STED confocal microscope (Leica TCS SP8 STED 3X). Images (Figure S2C) were acquired under 63× objective on Lattice SIM microscope (Zeiss Elyra 7) with wide-field mode. Confocal images (Figure 1H) were acquired under 63× objective on confocal microscope (Zeiss LSM 980 Airyscan2).

### 4.6. Live imaging

293T cells transfected with hCTPS1-mCherry and hCTPS1-miRFP670nano constructs were cultured on glass bottom culture dishes (C8-1.5H-N, Cellvis) with medium and maintained at 37 °C when live imaging was performed.

### 4.7. Image analysis

Fluorescence images was analyzed with the software IMAGEJ (NIH, Bethesda, MD, USA). The ROIs of bleached region for intensity measurement in Figure 2 were selected and measured manually with IMAGEJ. The FRAP curve unpaired t test was analyzed with Graph Prism 8. The deconvolution of STED images shown in Figure 3 was analyzed by lighting algorithm of Leica LAS X. The 3D model in Figure 3 was analyzed by Leica LAS X. The length of tentacles in figure 4 was measured manually with IMAGEJ. Quantity data was collected with Microsoft Excel.

### 4.8. Flow cell analysis

The flow cell analyzer used was LSRFortessa X20 (BD). Flow cell data was analyzed with FlowJo 10.4 software. Data was collected with Microsoft Excel.

## Author Contributions

Y. Fang and J-L. Liu conceived the project. Y. Fang designed experiments. Y. Fang, Y-L. Li and X. Li performed experiments. Y. Fang analyzed the data. Y. Fang wrote the original manuscript. Y. Fang, Y-L. Li, X. Li and J-L. Liu revised the manuscript.

## Acknowledgments

We thank the Molecular Imaging Core Facility (MICF) at the School of Life Science and Technology, ShanghaiTech University for providing technical support. This work was supported by grants from Ministry of Science and Technology of China (No. 2021YFA0804700), National Natural Science Foundation of China (No. 31771490), Shanghai Science and Technology Commission (No. 20JC1410500) and the UK Medical Research Council (grant nos. MC_UU_12021/3 and MC_U137788471) for grants to J.L.L.

## Data Availability Statement

Not applicable.

