## Supplemental Materials for "Super-resolution imaging reveals dynamic reticular cytoophidia"

### SUPPLEMENTAL INFORMATION

#### SUPPLEMENTAL FIGURES S1 AND S2

#### SUPPLEMENTAL TABLES S1 AND S2

### SUPPLEMENTAL FIGURES S1 AND S2 AND FIGURE LEGENDS

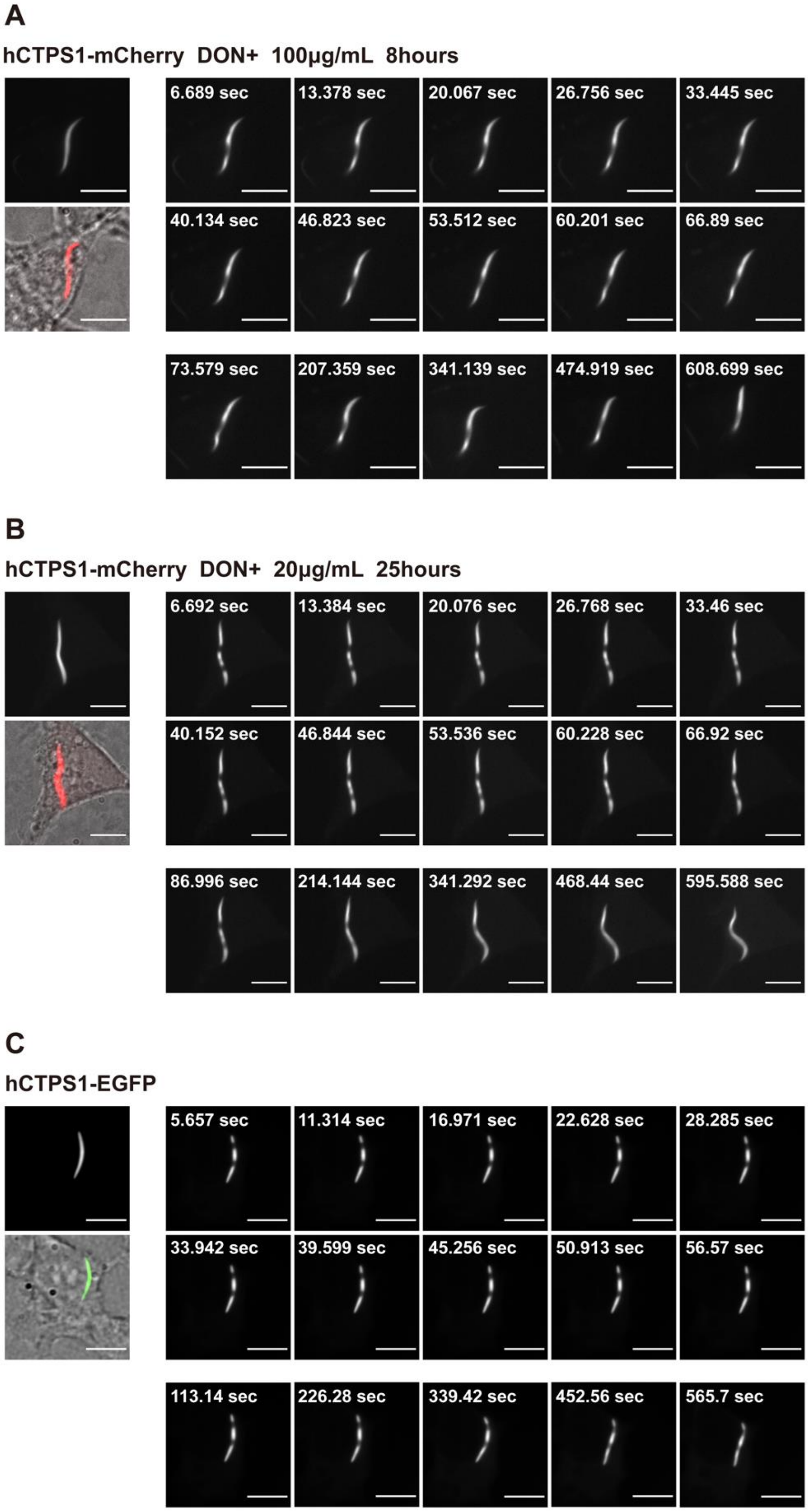

**Figure S1. FRAP on cytoophidia.**

(A) Living cell images of FRAP on hCTPS-mCherry cytoophidia induced by 100 μg/mL DON for 8 hours. (B) Living cell images of FRAP on hCTPS-mCherry cytoophidia induced by 20 μg/mL DON for 25 hours. (C) Living cell images of FRAP on hCTPS1-EGFP Cytoophidium-like structures. Scale bars, 10 μm (A-C).

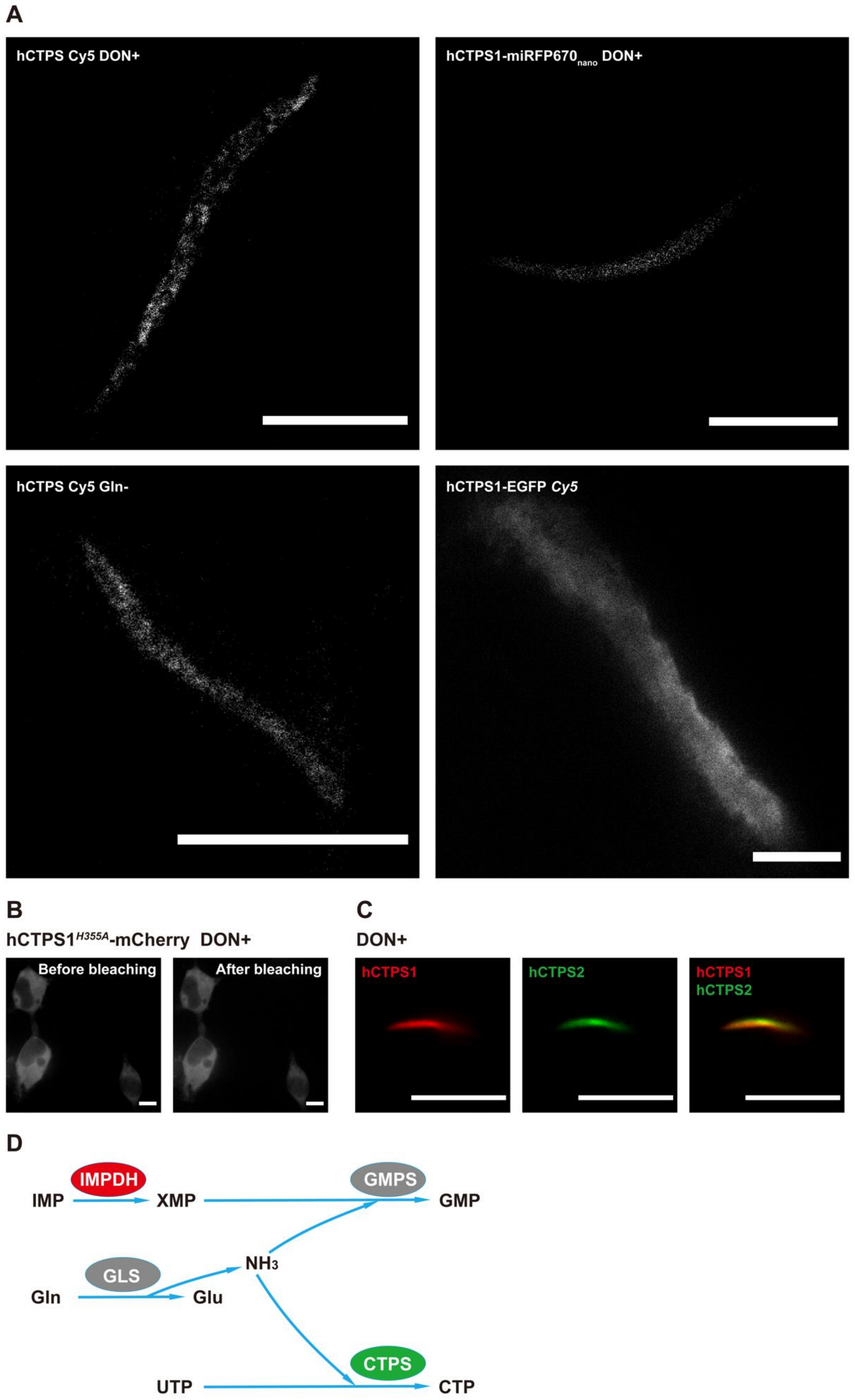
A
hCTPS Cy5 DON+
hCTPS1-miRFP670nano DON+
hCTPS Cy5 Gln-
hCTPS1-EGFP Cy5
B
hCTPS1H355A-mCherry DON+
Before bleaching
After bleaching
C
DON+
hCTPS1
hCTPS2
hCTPS1
hCTPS2
D
IMPDH
IMP
XMP
GMPS
GMP
NH3
GLS
Gln
Glu
CTPS
UTP
CTP

22

**Figure S2. Cytoophidia in super-resolution.**

(A) STED images before deconvolution of 1) hCTPS cytoophidia induced by DON in 293T cells, 2) hCTPS1 cytoophidia with miRFP670nano tags induced by DON in living 293T cells, 3) hCTPS cytoophidia induced by glutamine deprivation in SW480 cells and 4) hCTPS1-EGFP$^{K206A}$ cytoophidium-like structures in 293T cells. (B) There was no difference in hCTPS1 signal before and after bleaching. (C) hCTPS1 and hCTPS2 were colocalized under DON treated in the form of cytoophidia. For DON treatment, DON (PBS solution) was added into fresh DMEM media 8 hours before living cell imaging. Images are not analyzed by SIM algorithm. (D) IMPDH and CTPS are both parts of $NH_3$ metabolic pathways. Scale bars, 3 μm (A) and 10 μm (B, C).

#### SUPPLEMENTAL TABLES S1 AND S2

**Table S1. Over-expression proteins with fluorescence tags used in this study.**

| Exogenous proteins with tags | Figure |
|---|---|
| hCTPS1-mCherry | Figure 1A, B; Figure 2C; Figure 4A, C; Figure S1A, B; Figure S2B. |
| hCTPS1-EGFP$^{A206K}$ | Figure 1H, Figure 3C, D. |
| hCTPS1-EGFP | Figure 1H; Figure S1C. |
| hCTPS1$^{H355A}$-EGFP | Figure 1H. |
| hCTPS1-miRFP670nano | Figure 3A, Figure S2A. |
| hCTPS2-miRFP670nano | Figure S2C. |

**Table S2. List of oligonucleotides used in this study.**

| Primer name | Sequence (5'-3') | Target | Source |
|---|---|---|---|
| FYF-14F | ACGCGTTAAGTCGACAATCAACCTC | pLV-hCTPS1 | This paper |
| FYF-14R | GTCATGATTTATTGATGGAAACTTCAGTTCGGTG | pLV-hCTPS1 | This paper |
| FYF18-mCherry-F | TAAATCATGACCCACCGGTCATGGTGAGCAAGGGCGAGG | mCherry | This paper |
| FYF-13R | TGATTGTCGACTTAACGCGTTTACTTGTACAGCTCGTCCATGCCG | mCherry | This paper |
| FYF69-gfpK206A-F | GCCCTGAGCAAAGACCCCAACGAGAAGC | EGFP$^{K206A}$ | This paper |
| FYF70-gfpK206A-R | GTCTTTGCTCAGGGCGGACTGGGTGCTCAGGTAGTGG | EGFP$^{K206A}$ | This paper |
| FYF31F | GCAGAAGCTTGGCAGAAGCTCTGTAGTGC | hCTPS1$^{H355A}$ | This paper |
| FYF32-R | CTGCCAAGCTTCTGCGTAGCGCACGGGCTCTTC | hCTPS1$^{H355A}$ | This paper |
| FYF25-miRFPnano-FS | TAAATCATGACCCACCGGTCATGGCAAACCTGGACAAGATGCT | miRFP670nano | This paper |

| | | | |
|---|---|---|---|
| | GAA | | |
| FYF28-miRFPnano-RS | TTGTCGACTTAACGCGTTTAGCTCTGCTGGATGGCGATGC | miRFP670nano | This paper |
| FYF50-S2-temp-R | CAGGATGTACTTCATGGTGGCAGCGCTCTAGAACCG | miRFP670nano-pLV | This paper |
| FYF51-S2-temp-F | GAGTTGGAAATAAGCCCACCGGTCATGGCAAACCTGG | miRFP670nano-pLV | This paper |
| FYF43-hCTPS2-F | ATGAAGTACATCCTGGTCACGGGTGG | hCTPS2 | This paper |
| FYF44-hCTPS2-R | GCTTATTTCCAACTCAGCTATCCTTGGCTCT | hCTPS2 | This paper |
